# Ants express risk-adjusted sanitary care

**DOI:** 10.1101/170365

**Authors:** Matthias Konrad, Christopher D. Pull, Katharina Seif, Sina Metzler, Anna V. Grasse, Sylvia Cremer

**Affiliations:** IST Austria (Institute of Science and Technology Austria), Am Campus 1, A-3400 Klosterneuburg, Austria.

**Author notes:** Corresponding author: Sylvia Cremer. Equal contribution (Matthias Konrad, Christopher D. Pull). Current address: Medical University of Vienna, Schwarzspanierstr. 17, A-1090 Vienna, Austria (Katharina Seif).

**Keywords:** host-pathogen interactions, social immunity, behavioural plasticity, co-infection

## Abstract

Being cared for when sick is a benefit of sociality that can reduce disease and improve survival of group members. However, individuals providing care risk contracting infectious diseases themselves. If they contract a low pathogen dose, they may develop micro-infections that do not cause disease, but still affect host immunity by either decreasing or increasing the host’s vulnerability to subsequent pathogen infections. Caring for contagious individuals can thus significantly alter the future disease susceptibility of caregivers. Using ants and their fungal pathogens as a model system, we here tested if the altered disease susceptibility of experienced caregivers, in turn, affects their expression of sanitary care behaviour. We found that micro-infections contracted during sanitary care had protective or neutral effects upon secondary exposure to the same (homologous) pathogen, but consistently induced high mortality upon super-infection with a different (heterologous) pathogen. In response to this risk, the ants selectively adjusted the expression of their sanitary care. Specifically, the ants performed less grooming yet more antimicrobial disinfection, when caring for nestmates contaminated with heterologous pathogens as compared to homologous ones. By modulating the components of sanitary care in this way, the ants reduced their probability of contracting super-infections of the harmful heterologous pathogens. The performance of risk-adjusted sanitary care reveals the remarkable capacity of ants to react to changes in their disease susceptibility, according to their own infection history, and to flexibly adjust collective care to individual risk.

## Introduction

The infection history of a host can severely impact its future disease susceptibility. For example, previous infections or vaccinations can immunise and thus protect hosts against secondary infections of the same, homologous pathogen [1, 2]. Yet, prior infection can also increase a host’s susceptibility to other pathogens, leading to heterologous super-infections that have detrimental impacts on host health and survival [3, 4]. In addition to immunity, behaviour often also changes upon infection: on the one hand, pathogens can manipulate host behaviour to facilitate disease transmission [5]; on the other, changes in host responses – termed sickness behaviour – can speed up recovery and limit pathogen spread, e.g. through reduced activity and modulation of social interactions [6-8]. Infection can even alter an animal’s sensitivity to pathogen-associated stimuli, as, for example, disease avoidance behaviour [“disgust”; 9, 10] can be affected by a host’s infection history [11, 12].

Disease prevention is particularly important in social groups, where pathogens can easily spread due to a high density of hosts and the multitude of interactions between them [13-15]. Consequently, social animals have evolved behavioural disease defences, which operate in conjunction with individual immunity, to negate the increased pathogen risk they experience [16-18]. For example, social insects have evolved sophisticated, collective defences against diseases that result in an emergent, group-level protection of the colony, known as social immunity [18]. One particularly important component of social immunity is sanitary care – grooming to remove pathogens [19, 20] and chemical disinfection to inhibit their growth [21, 22] – that effectively reduces the risk of disease for pathogen-exposed individuals. However, this intimate behavioural interaction often also involves pathogen transmission from the contaminated individual to those performing sanitary care. Insects caring for their nestmates can therefore either become sick themselves [19, 20], or contract micro-infections that do not lead to disease symptoms, but can confer protection against the same, homologous pathogen, upon secondary exposure. For example, in both ants [23] and termites [24], social contact with contaminated individuals carrying infectious conidiospores of the entomopathogenic fungus *Metarhizium* on their cuticle causes micro-infections in most of their nestmates. Interestingly, these micro-infections trigger an upregulation of antifungal genes, and hence antifungal activity of the insects [23, 24], which leads to a significant survival benefit – known as social immunisation [2] – upon secondary challenge with the same, homologous pathogen [25, 26]. In humans, prior natural infections and vaccination affect individual pathogen avoidance behaviour [12] and strategies for providing health care [27, 28]. However, in social insects, it is unknown how infection history affects future host susceptibility to different (heterologous) pathogens, and whether micro-infections can alter the performance of sanitary care of ants.

To address this, we set up a full-factorial experiment, using the natural host-pathogen system of garden ants and entomopathogenic fungi (hosts: *Lasius* ants; pathogens: *Metarhizium Beauveria*) [29, 30]. These two pathogens have similar reproductive cycles: infectious conidiospores are acquired from the environment/infectious corpses and initially attach loosely to an insect’s cuticle [31, 32]. At this time, conidiospores can be removed from contaminated ants through grooming [20, 33], or inactivated by disinfection to prevent infection [21, 22]; additionally, at this stage, conidiospores can be transferred from contaminated individuals to those performing sanitary care [23]. However, once the conidiospores fully adhere to the cuticle, they can no longer be removed [34] and will penetrate into the hemocoel of the insect, causing an internal infection [35]. If the infective dose is high enough to overcome the immune system, the fungus will kill the host and grow out of the body, eventually producing new conidiospores on the corpse [20, 31].

To elucidate the effects of micro-infections acquired through sanitary care on future individual disease susceptibility and sanitary care in ants, we induced micro-infections in ants by rearing *Lasius neglectus* workers with a pathogen-exposed individual, treated with either *Metarhizium robertsii* or *Beauveria bassiana*, for five days [as in 23]. We set up an additional non-infected control group by keeping workers with a sham-treated individual, which could not transfer any pathogen. We then tested how acquired micro-infections affect ant mortality upon a challenge with the same or the heterologous pathogen, whether micro-infections change the expression of sanitary care, and if these changes alter disease transmission.

## Results and Discussion

### Micro-infections increase an ant’s susceptibility to heterologous super-infection

We compared the effect of pathogen challenge in ants with micro-infections of either a homologous (*Metarhizium–Metarhizium, Beauveria–Beauveria*), or a heterologous (*Metarhizium–Beauveria, Beauveria–Metarhizium*) pathogen. Control ants had no previous micro-infection (non-infected–*Metarhizium,* non-infected–*Beauveria*). Ant mortality significantly differed between the groups (Fig. 1; Cox mixed effects model: likelihood ratio test (LR) χ^2^ = 11.31, df = 2, *P* = 0.004). Yet, as micro-infections homologous to the subsequent challenge only provided a protective effect for *Metarhizium*, but not for *Beauveria* (Fig. S1 and statistics therein), we did not find an overall survival benefit of a homologous micro-infection compared to the non-infected controls (post hoc comparisons: non-infected vs. homologous, *P* = 0.17, hazard ratio (HR) = 0.79). Our data thus reveals that, in addition to *M. brunneum* [23], protective social immunisation in garden ants can also be elicited by other *Metarhizium* species. However, the neutral effect for *B. bassiana* micro-infections shows that social immunisation is induced by some, but not all, pathogens, similar to individual and transgenerational immune priming of invertebrates [reviewed in 2].

**Fig. 1).**
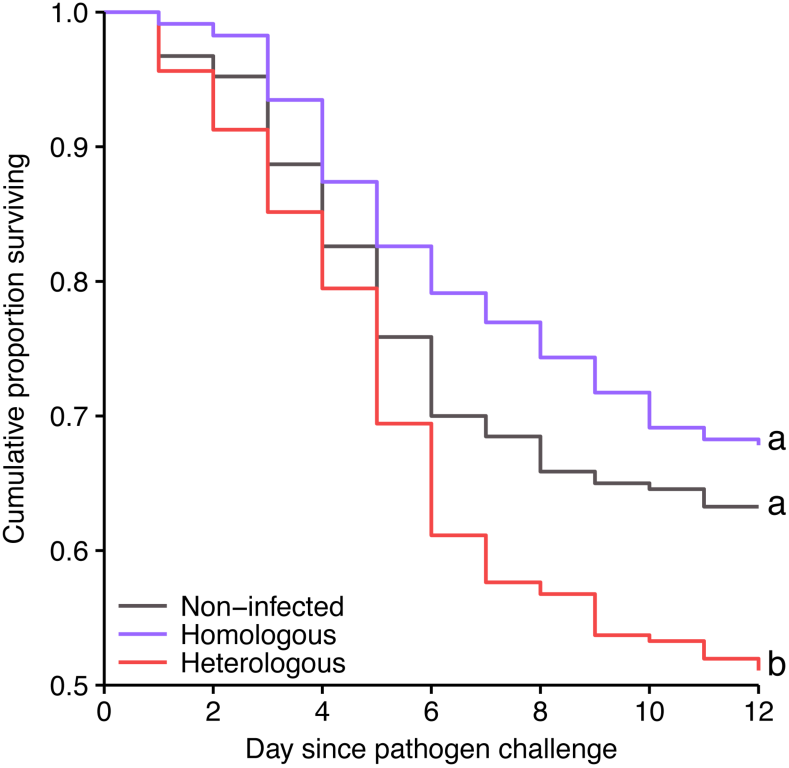
Increased mortality of micro-infected ants after heterologous pathogen challenge. Pathogen exposure induced significantly higher mortality in *Lasius neglectus* ants that had an existing micro-infection with a heterologous pathogen (red line), compared to both, a micro-infection with the homologous pathogen (purple) or no infection (grey), with the latter groups not differing significantly from one another. Different letters indicate significance groups of all pairwise post-hoc comparisons after Benjamini-Hochberg correction at α=0.05. For separate analysis of the two pathogens (*Metarhizium* and *Beauveria*) see Fig. S1.

Heterologous combinations of micro-infection and challenge with a pathogen from a different fungal genus (*Metarhizium–Beauveria, Beauveria–Metarhizium*) invariably increased ant mortality, both compared to the non-infected controls (*P* = 0.024, HR = 1.47) and the homologous challenge (*P* = 0.003, HR = 1.86), which was independent of pathogen order (Fig. S1). To test if this effect was dependent on whether one of the two pathogens had already established an infection in the host body, or if it reflects a general pattern of *Beauveria–Metarhizium* co-infection, we simultaneously co-exposed ants to a mix of both pathogens, keeping the overall pathogen load constant. Again, we found a strong mortality-inducing effect compared to ants exposed to only the single pathogens (Cox proportional hazards regression: LR χ^2^ = 45.1, df = 1, *P* = 0.001, HR = 2.68; independent of pathogen species; Fig. S2 and statistics therein). Similar harmful effects are well-established for heterologous challenges after individual and transgenerational immune priming [2, 36, 37] and concurrent infections with different pathogens [3, 4]. They can result either from pathogen-pathogen interactions, including both competition and cooperation [38, 39], or from perturbations of the host immune system [40, 41].

### Micro-infected ants modulate their sanitary care

Social insect colonies face a high load [42] and diversity [43, 44] of pathogens in their environment, making multiple infections of individual colony members likely. Given the observed costs of increased susceptibility to heterologous pathogens after a previous, otherwise asymptomatic micro-infection, we expect selection to act on ants to avoid super-infections with detrimental heterologous pathogens. As contaminated individuals represent a constant risk of infection for their colony members [19, 20, 23], we tested the hypothesis that ants should modulate their behaviour to selectively reduce the risk of heterologous pathogen contraction when in contact with a pathogen- contaminated nestmate. To this end, we confronted micro-infected ants with a contaminated nestmate, from which they could contract a super-infection of either the homologous or heterologous pathogen, and observed how their behaviour differed from that of non-infected ants (Movie S1).

While aggression between nestmates is usually absent in *L. neglectus* colonies [45], micro-infected ants started biting, grabbing and dragging their contaminated nestmates. This was equally so, whether or not the nestmate was contaminated with the homologous or heterologous pathogen to their micro-infection (Fig. 2A; generalised linear model (GLM) χ^2^ = 13.39, df = 2, *P* = 0.002; post hoc comparisons: non-infected vs. homologous, *P* = 0.007; non-infected vs. heterologous, *P* = 0.0002; homologous vs. heterologous, *P* = 0.24; see Figs. S3A,B, table S1 for separate statistical analyses for the two pathogen species, showing a significant effect for nestmates contaminated with *Metarhizium* and the same, yet non-significant pattern in *Beauveria*). Hence, whilst increased aggression has previously been reported for infected ants towards non-nestmates from different colonies [46], and by healthy ants and honeybees towards infected or immune-challenged nestmates [47-49], here we find that micro-infected ants have a heightened level of aggression themselves, towards their own nestmates.

**Fig. 2).**
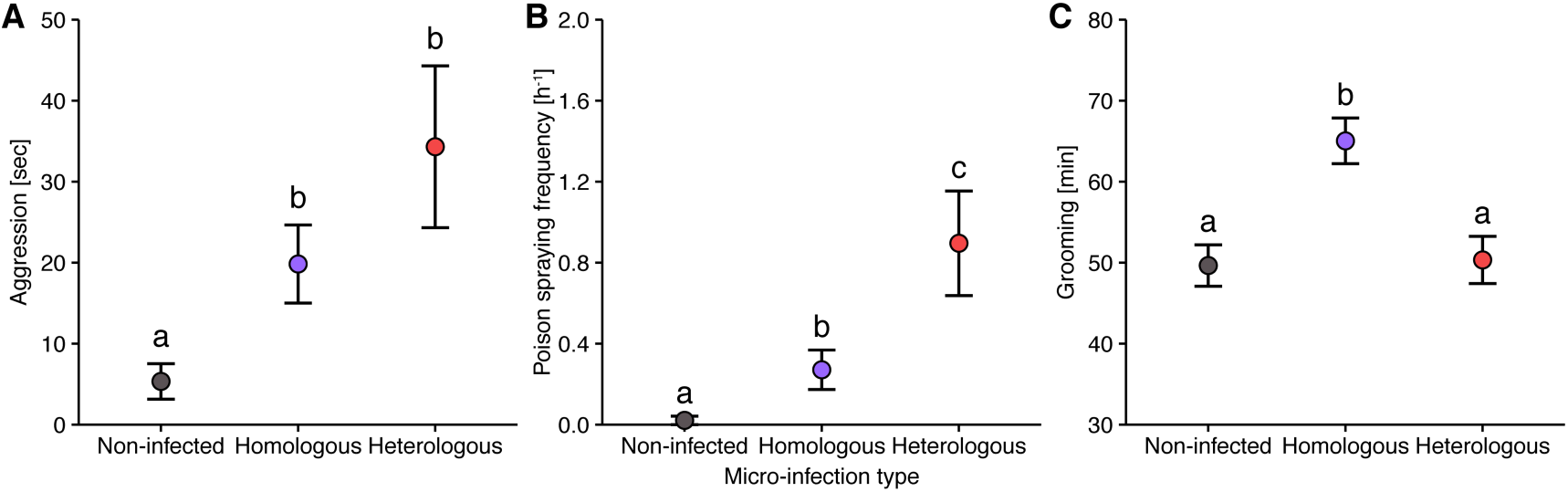
Modulation of behaviour displayed by micro-infected ants towards contaminated nestmates. Micro-infected ants differed in their aggression and sanitary care behaviours from non-infected control ants (grey symbols), and displayed behavioural plasticity depending on whether their nestmate was contaminated with the homologous (purple) or heterologous pathogen (red) to their micro-infection. (A) Micro-infected ants became significantly more aggressive towards their nestmate, irrespective of whether it was contaminated with the homologous or heterologous pathogen. (B) Poison spraying was essentially absent in non-infected control ants, but significantly increased in micro-infected ants interacting with a nestmate contaminated with the homologous pathogen, and further significantly increased when the nestmate was contaminated with the heterologous pathogen. (C) Grooming was performed significantly longer by micro-infected encountering a nestmate contaminated with the homologous pathogen, compared to both non-infected control ants and micro-infected ants grooming nestmates contaminated with heterologous pathogen. Mean ± SEM displayed; different letters indicate significance groups of all pairwise post-hoc comparisons after Benjamini-Hochberg correction at α=0.05. For separate analysis of the two pathogens (*Metarhizium* and *Beauveria*) see Fig. S3.

Despite increased aggression, micro-infected ants still performed sanitary care – grooming and chemical disinfection [21] – towards their contaminated nestmates. When grooming, ants remove infectious particles with their mouthparts from the cuticle of nestmates [20, 33, 50]. In addition, *L. neglectus* ants spray their formic acid rich, antimicrobial poison onto exposed colony members – an effective disinfection behaviour that is distinct from defensive poison use [21]. The application process itself is a fast and infrequent behaviour, followed by a long replenishment phase of the poison reservoir [21]. We found that the relative expression of grooming and poison spraying in micro-infected ants was dependent on whether the nestmate was contaminated with the homologous or heterologous pathogen (Fig. 2B,C).

Micro-infected ants sprayed poison towards their contaminated individuals, whilst this behaviour was essentially absent in non-infected ants (Fig. 2B; GLM, χ^2^ = 21.88, df = 2, *P* = 0.0002; post hoc comparisons: non-infected vs. homologous, *P* = 0.021; non-infected vs. heterologous, *P* = 0.002). Micro-infected ants sprayed poison significantly more often towards nestmates contaminated with the heterologous, than the homologous pathogen (Fig. 2B; *P* = 0.021; whilst these effects were only a non-significant trend at *P* = 0.08 for nestmates contaminated with *Metarhizium*, and non-significant for nestmates contaminated with *Beauveria* alone; see Figs. S3C,D and table S1 for separate statistical analyses for the two pathogen species).

Ants also expressed differences in allogrooming depending on their micro-infection and pathogen threat (Fig. 2C; GLM, *F* = 9.87, df = 2, *P* = 0.0002). As observed in other species [34, 50], prior pathogen encounter caused an increase in the grooming of nestmates contaminated with a homologous pathogen (post hoc comparisons: non-infected vs. homologous, *P* = 0.0003). Importantly, however, we found that grooming did not increase when ants encounter nestmates contaminated with a heterologous pathogen (non-infected vs. heterologous, *P* = 0.86) and was hence significantly more frequent in interactions with nestmates carrying the homologous than the heterologous pathogen (*P* = 0.003; which was significant for nestmates contaminated with *Metarhizium* and showed the same, yet non-significant, pattern for *Beauveria*; see Figs. S3E,F and table S1 for separate analyses for the two pathogen species).

Overall, our behavioural data show that micro-infected ants start becoming aggressive towards their contaminated nestmates, but nevertheless still perform sanitary care. However, they modulate the components of sanitary care, in that micro-infected ants perform more poison spraying but less grooming of nestmates contaminated with a heterologous, compared to the homologous pathogen. Since grooming is a more intimate behaviour than poison spraying, it is likely to lead to more pathogen transmission between nestmates. By modulating their behaviour in this way, micro-infected ants may therefore minimise the risk of transmission when nestmates are contaminated with a harmful heterologous pathogen, to which they are more susceptible.

### Risk-averse sanitary care reduces heterologous pathogen transfer and infection load

To determine if the observed behavioural modulation of micro-infected ants affects disease transmission, we first tested if micro-infected ants contracted less of the detrimental heterologous pathogen, and, secondly, if this had a beneficial effect on the ants’ infection load and, thus, risk of disease.

To simultaneously measure pathogen transmission and fungal growth inhibition by antimicrobial poison spraying, we determined the number of infectious fungal conidiospores that were transferred to the body surface of ants performing sanitary care, immediately after their social interaction with contaminated nestmates. We found that micro-infected ants acquired significantly less infectious particles while interacting with nestmates contaminated with the heterologous pathogen, as compared to both micro-infected ants interacting with nestmates contaminated with the homologous pathogen, or non-infected ants from contaminated nestmates, with the latter two not differing from one another (Fig. 3A; GLM, *F* = 8.79, df = 2, *P* = 0.0003; post hoc comparisons: non-infected vs. homologous, *P* = 0.85; non-infected vs. heterologous, *P* = 0.0008; homologous vs. heterologous, *P* = 0.0008). These results were robust, independent of pathogen order (see Fig. S4 and statistics therein). This reveals that the behavioural plasticity displayed by the ants depending on the combination of their micro-infection and pathogen threat, reduced the transfer of the infectious heterologous, but not of the non-detrimental homologous, pathogen compared to non-infected control ants.

**Fig. 3).**
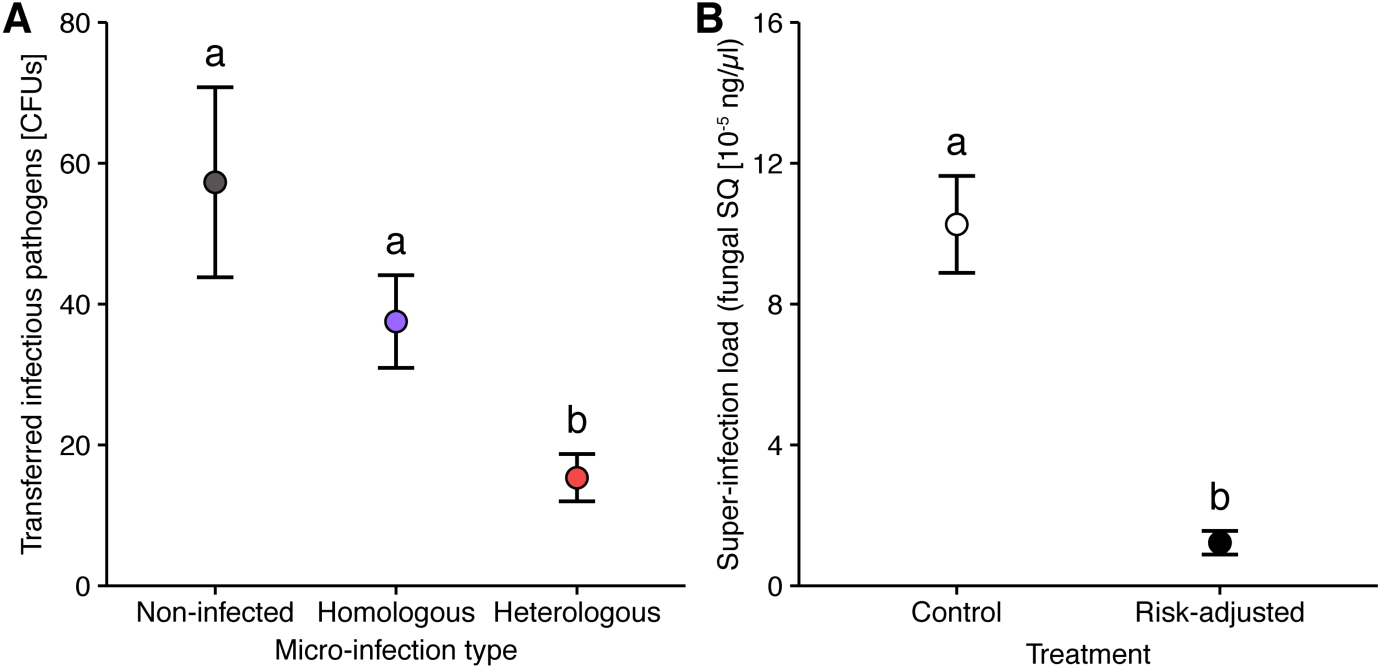
Reduced heterologous pathogen transfer and super-infection load to micro-infected ants. (A) Micro-infected ants acquired significantly lower amounts of viable, infectious pathogen during social contact with nestmates contaminated with the heterologous pathogen (red symbol) than both, micro-infected ants interacting with nestmates contaminated with the homologous pathogen (purple), or non-infected controls interacting with contaminated nestmates (grey), with the latter two groups not differing significantly. (B) We measured the super-infection levels of the heterologous pathogen when micro-infected ants were exposed to fungal conidiospore dosages equivalent to what they naturally acquired during interactions with a nestmate contaminated with the heterologous pathogen (Fig. 3A heterologous; filled symbol), as compared to control ants (Fig. 3A controls, open symbol). We found that super-infection levels were significantly lower in ants displaying risk-averse behavioural modulation compared to the control ants. Mean ± SEM displayed; different letters indicate significance groups of all pairwise post-hoc comparisons after Benjamini-Hochberg correction at α=0.05. For separate analysis of the two pathogens (*Metarhizium* and *Beauveria*) see Figs. S4 and S5.

Since exposure dose is related to infection load and the risk of disease [51], we tested whether the reduced transfer of harmful heterologous pathogen translates into a lower super-infection load in micro-infected ants. To this end, we exposed micro-infected ants to two different dosages of their heterologous pathogens: (i) a low dose equivalent to the amount ants received when performing risk-adjusted sanitary care (determined from data in Fig. 3A - heterologous pathogen combinations) or (ii) a higher dose equivalent to the amount received when the behavioural change was absent (determined from data in Fig. 3A – non-infected control ants). Our data clearly show that the reduced transfer of infectious conidiospores in ants displaying risk-adjusted sanitary care minimises their super-infection load (Fig. 3B; linear mixed effects regression (LMER), χ^2^ = 41.44, df = 1, *P* = 0.0001; independent of pathogen order; Fig. S5 and statistics therein). Since the disease caused by the pathogens we used is dose-dependent [51], we can thus conclude that the risk-adjusted sanitary care displayed by micro-infected ants will reduce their chances of contracting the heterologous pathogen (to which they are more susceptible) and developing a detrimental super-infection.

## Conclusion

In this study, we investigated how micro-infections acquired through sanitary care affect an ant’s disease susceptibility to future infections with the same, homologous or a different, heterologous pathogen, and, as a consequence, how this affects their interactions with pathogen-exposed nestmates. We found that micro-infections had a protective or neutral effect when ants were exposed to the homologous pathogen, but increased an ant’s susceptibility to super-infections of heterologous pathogens (Fig. 1). However, by altering the relative expression of grooming and poison spraying (Fig. 2), micro-infected ants were able to specifically reduce the amount of heterologous pathogens they contract when performing sanitary care of contaminated nestmates (Fig. 3A). Importantly, this risk-averse behaviour results in a lower infection load for micro-infected ants (Fig. 3B), meaning their chances of acquiring super-infection with a detrimental pathogen are reduced.

Micro-infected ants were more aggressive to their nestmates, indicating that the infection itself or a physiological change due to an immune response alters the level of aggression in ants. Similar to our findings in ants, higher irritable or hostile behaviour can also be observed in infected vertebrates, including humans [52, 53]. These changes in behaviour form part of a general sickness behaviour that also includes fatigue, depression, and reduced social integration [6-8]. Such changes in behaviour are generally considered to be triggered by the immune system through neural and circulatory pathways [54]. In ants, a loss of attraction to social cues from the nest or nestmates [55], and hence social isolation from the colony, occurs after *Metarhizium* infection, close to the death of the individual [46, 47]. Yet, the social isolation of diseased, or generally moribund [56], ants does not involve any aggressive interactions in the colony [47, 55]. The observed increase in aggression levels in micro-infected ants may thus constitute a “partial sickness behaviour”, as they neither express signs of disease nor all components of a typical sickness behaviour [6, 55], and still actively engage in sanitary care of their contaminated nestmates. Whether these aggressive behaviours are adaptive remains to be determined, but dragging and biting of infected individuals has been observed in honeybees and other ants [47-49], and may potentially play a role in removing contagious individuals from the colony, to reduce disease spread [6].

Micro-infected ants continued to provide sanitary care to their contaminated nestmates, but in a risk-averse manner. Remarkably, we found that by altering the expression of sanitary care, ants are able to reduce their own risk of contracting a super-infection with a second, harmful pathogen, whilst caring for their contaminated nestmates. This self-protection is achieved by adjusting the relative expression of the components of sanitary care – grooming vs. disinfection. Specifically, micro-infected ants intensified the spraying of their antimicrobial poison [21] but did not increase the mechanical removal of the infectious particles by grooming of the body surface of nestmates [20], when they were contaminated with the heterologous pathogen. As grooming involves close contact and uptake of the infectious particles into the ants’ infrabuccal pockets within their mouths [21, 57], whilst poison spraying represents only the application of disinfectant, this shift in sanitary care led to a reduction in pathogen transfer. The capacity to specifically prevent super-infections of detrimental, heterologous pathogens is expected to benefit both the host and the pathogen that has already established a micro-infection [58]. Consequently, elucidating the underlying mechanistic basis of the behavioural changes in micro- infected ants will be necessary to determine the relative importance of an adaptive shift in the host’s response and potential parasite manipulation [59].

Importantly, ants did not simply perform risk-averse sanitary care against pathogens they inherently were more or less susceptible to, e.g. due to their genetic predisposition. Instead, the ants reacted to their individual infection history that induced a *change* in their disease susceptibility. Our data hence represent a prime example of behavioural plasticity, showing how sanitary care is adjusted as a consequence of the combination of the individual infection history of the care-providing insects, and the encountered second pathogen, present on the contaminated nestmate. Such behavioural modulation, based on susceptibility changes due to prior experience [60], is also known in flies [61] and humans, e.g. during pregnancy [11, 62]. However, in these cases, the changes lead to an increased avoidance of infectious individuals – a behaviour that occurs in healthy animals from crustaceans to primates [63-65] – rather than a modulation of care behaviour, as we report here. The ultimate outcome, the performance of risk-averse sanitary care, rather than the avoidance of diseased group members, highlights the importance of social immunity for the success of insect societies. Indeed, although workers are often considered to be expendable, they are essential for colony maintenance and the production of new queens, so their health status will have a direct impact on colony fitness [66]. Hence, behavioural adaptations, like those presented here, ensure that the workerforce of the colony remains healthy and, consequently, can contribute productively to enhance colony fitness.

## Methods

### Ant hosts

We used the invasive garden ant, *Lasius neglectus* [45, 67] sampled from Jena, Germany (N 50° 55.910 E 11° 35.140), and reared in the laboratory (as in [23]). Ants were kept at a constant temperature of 23°C with 75% humidity and a day/night cycle of 14 h light/10 h dark. Experiments were performed in petri dishes with a plastered base and 10% sucrose solution as an *ad libitum* food supply. Ants were randomly assigned to the respective treatment groups, described below. *L. neglectus* is an unprotected insect species, and all experiments comply with European laws and IST Austria ethical guidelines.

### Fungal pathogens

We used the entomopathogenic fungi *Metarhizium* and *Beauveria*, both of which are natural pathogens of *Lasius* ants [29; C.D.P unpubl., 30] and occur in high density (up to 5000 conidiospores/g soil, [42]) and diversity (many sympatrically occurring species and strains, [43, 44]) in the soil, where the ants nest. Natural infection loads of *L. neglectus* populations with individual species of these obligate killing pathogens reach up to 9% prevalence [29], with sporulating cadavers each producing approximately 12 Mio new infectious conidiospores (conidia) [20]. In *Metarhizium*, topical application of approximately 30 conidiospores induces 2% mortality in *L. neglectus* workers [23], whilst application of 300,000 spore constitutes the lethal dose (LD)50 [23]. We used the strains *Metarhizium robertsii* KVL 13-12 and *Beauveria bassiana* KVL 04-004 (obtained from the University of Copenhagen, Denmark), of which multiple aliquots were kept in long-term storage at −80°C. Prior to each experiment conidiospores of both fungi were grown on malt extract agar at 23°C for 3 w and harvested by suspending them in 0.05% sterile Triton X-100 (Sigma). All conidiospore suspensions had a germination rate of > 91%, which was determined directly before each experiment.

### Fungal exposure of ants

We exposed individual worker ants by applying a 0.3 µl droplet of conidiospore suspension onto their gaster (abdomen) and subsequently letting them dry for 1 min on filter paper. The concentration of each conidiospore suspension was adjusted depending on the respective fungus and experiment (see below). For the sham control treatment we applied a 0.3 µl droplet of 0.05% Triton X-100 (Sigma) solution only.

### Establishment of micro-infections

To induce socially-acquired micro-infections, we grouped 5 naive ants with a single pathogen-exposed individual (distinguishable by paint marking the exposed individual [Edding 780]) in a petri dish (Ø = 9 cm; as in [23]). The exposed individual was either treated with *Metarhizium* or *Beauveria* (both 1×10^9^ conidiospores/ml). To obtain non-infected control ants, 5 naive ants were grouped with a sham-treated individual. After 5 d of social contact the treated individual was removed and the remaining ants – micro-infected with either *Metarhizium* or *Beauveria*, or non-infected controls – were subjected to further experiments (see below).

### Survival of micro-infected ants upon homologous and heterologous pathogen challenge

Ants micro-infected with *Metarhizium* or *Beauveria*, as well as non-infected controls, were exposed to either *Metarhizium* (1×10^9^ conidiospores/ml) or *Beauveria* (5×10^9^ conidiospores/ml) and subsequently kept as single ants in new petri dishes (Ø = 3.5 cm). Survival of the ants was monitored daily for a period of 12 d. To confirm deaths caused by internal infection of the entomopathogenic fungi, ant corpses were surface-sterilized (using bleach and ethanol, [68]), kept under humid conditions for 3 w and regularly checked for fungal outgrowth and conidiospore formation. Our experimental setup thus comprised 4 different combinations of ‘micro-infection’ and ‘pathogen challenge’ (2 homologous and 2 heterologous), each with its own control group (‘non-infected ants’ + ‘pathogen challenge’). Each combination comprised 24 replicates of 5 micro-infected or control ants per petri dish, resulting in 120 ants per combination and 960 ants overall in the experiment. 4.3% of the ants (41/960) died before direct pathogen challenge and were therefore excluded from further statistical analyses. The ants’ infection history (i.e. non-infected *versus* micro-infected) did not affect mortality before pathogen challenge (Chi-square test: χ^2^ = 0, df = 1, *P* = 1). 83.1 % of dead ants (295/355) showed fungal outgrowth after surface sterilization, thus confirming an internal entomopathogenic infection.

### Simultaneous co-exposure with *Metarhizium* and *Beauveria*

We studied the effects of pathogen co-infection by exposing naive ants (n = 120 per treatment, total 360 ants) to a mixture of both fungal pathogens (each 50%; total 1×10^9^ conidiospores/ml), or each pathogen alone (as above). Exposed ants were reared singly (petri dishes: Ø = 3.5 cm) and survival was monitored daily for 12 d. Dead ants were surface sterilized (as above), with 89.6% (155/173) showing fungal outgrowth (89.1% for *Metarhizium-*, 91.3% for *Beauveria*- and 89.5% for co-exposure).

### Behavioural observations of micro-infected ants

We observed the behaviour of micro-infected ants towards a newly encountered contagious nestmate. To this end, groups of 5 ants micro-infected with either *Metarhizium* or *Beauveria,* or non-infected control ants, were transferred into petri dishes (Ø = 3.5 cm; no food). A colour-marked nestmate from their parental colony, with whom the ants had no contact to in the previous 5 d, was then introduced after exposure to either *Metarhizium* or *Beauveria* (both 1×10^9^ conidiospores/ml). This led to six combinations of micro-infection (non-infected, *Metarhizium*, or *Beauveria*) and nestmate contamination treatment (*Metarhizium*, or *Beauveria*), each consisting of 24 replicates (n = 5 micro-infected or non-infected control ants per petri dish, i.e. 120 per combination, 720 in total).

After an acclimatisation period of 10 min following nestmate introduction, we recorded the ants’ behaviour towards the treated nestmate for 1 h (camera: Di-Li 970-O; software: Debut video capture 1.64; always simultaneously filming one replicate each of the non-infected, *Metarhizium-*, or *Beauveria-* micro-infected treatment group with random assignment to the three cameras running in parallel). Videos were analysed “blind” with regard to treatment, using the software ‘Biologic’ (http://sourceforge.net/projects/biologic/), to record the start and end time point of the ants’ behaviour. We then calculated (i) the total duration (in sec) of aggression (comprising aggressive picking/licking/plucking of the body surface, mandible biting into different body parts, and dragging), (ii) the number of events of direct poison spraying from the acidopore (opening of the poison gland at the gaster tip), as well as (iii) the total duration of allo-grooming (in sec) that the treated nestmate received, per replicate (see Movie S1 for examples).

### Conidiospore transfer from the contaminated nestmate to micro-infected ants

To study if the ants’ infection history affected their likelihood of acquiring the homologous *vs.* heterologous pathogen when interacting with a contaminated nestmate, we determined the number of viable colony forming units (CFUs) of conidiospores on the body surface of non-infected and micro-infected ants (*Metarhizium*, *Beauveria*), after 70 min of social contact to their *Metarhizium-* or *Beauveria-*contaminated nestmate, again in all possible combinations (same experimental setup as above). Ants (total n = 820 individuals, pooled in 164 replicates of 5 ants each) were transferred into tubes (1.5 ml) containing 100 µl Triton X (0.05 %), to wash off and collect conidiospores that these ants may have acquired from their contaminated nestmate. The tubes were shaken for 10 min on a vortex mixer (Vortex-Genie 2, Scientific Industries) at maximum speed and ants were subsequently removed from the tubes. We plated the washes on selective medium agar plates (containing: chloramphenicol 100 mg/l, streptomycin 50 mg/l, dodin 110 mg/l), cultivated the plates at 23°C for 2 w and determined the number of colony forming units (CFUs) of *Metarhizium* and *Beauveria* growing on each plate through visual inspection.

We confirmed that this procedure is appropriate to quantify only the conidiospore transfer during contact with the contaminated nestmates, but not any conidiospores remaining from the previous micro-infection induction five days prior. To this end, we let non-infected or micro-infected (*Metarhizium*, or *Beauveria*) ants interact with nestmates that had only received a sham-treatment and thus could not transfer pathogen (10 replicates of 5 ants each; total 150 ants). None of these washes led to growth of entomopathogenic fungi after 2 w of cultivation, revealing that the micro-infected ants did not carry any viable conidiospores on their cuticle from their previous micro-infection that could still be washed off at this time point. Consequently, all viable conidiospores from our washes can only have been transferred from the contaminated individual within the 70 min of social contact in our experiment.

### Relationship between conidiospore transfer and internal fungal load of ants

We determined whether the observed differences in conidiospore transfer between ants displaying the behavioural change, as compared to ants that do not, translates into different levels of super-infection, which establish in the ants after exposure. To this end, we directly exposed micro-infected ants to the heterologous pathogen, in an amount they would acquire when expressing the behavioural change (i.e. micro-infected ants interacting with a nestmate contaminated with the heterologous pathogen), and compared their super-infection load with that of micro-infected ants, where we simulated an absence of the behavioural change by exposing them to the heterologous pathogen in an amount non-infected control ants, which do not express the behavioural change, would acquire (Fig. 3A, Fig. S4).

As the accuracy to determine conidiospore transfer by washing (see above) diminishes over time due to increasing conidiospore attachment, yet transfer between ants is possible for approx. 48 h after exposure in this experimental system [34; own obs.], we had to determine the required application doses by extrapolation from our early observation period (first 70 min of interaction) to the full period of possible pathogen transfer. Within the 48 h of possible transmission, the likelihood of conidiospore transfer continuously decreases with time, firstly, because a constantly reducing number of conidiospores can be transferred per interaction due to reduction in number via grooming, or stronger attachment to the cuticle due to germination, and, secondly, because the ants perform fewer grooming interactions per unit of time. In our system, grooming decreases by 40% between 24 and 48 h after exposure [23]. We therefore estimated how many conidiospores are transferred within the full infective period of 48 h using an exponential function, taking the number of conidiospores transferred within the first hour (Fig. 3A) as the starting point and a decay rate *tau* of 10, based on our observations in [23]. As experimental topical application further is a ‘wasteful procedure’ and only 10-15% of the applied conidiospores are in fact sticking to the body surface (MK & SM unpubl.), we used the following application dosages in our experiment: *Beauveria* micro-infected: behavioural change present = 460 *Metarhizium* conidiospores, behavioural change absent =1850; *Metarhizium* micro-infected: behavioural change present = 90 *Beauveria* conidiospores, behavioural change absent = 1480.

After having been kept with a contaminated individual for 5 d to acquire a micro-infection (as above), ants were directly exposed to the heterologous fungus and kept in isolation (petri dishes: Ø = 3.5 cm) for another 5 d. After this time ants were freeze-killed and later combined into pools of 5 individuals each (number of replicates, *Beauveria* micro-infected: behavioural change present = 14, behavioural change absent =13; *Metarhizium* micro-infected: behavioural change present = 12, behavioural change absent = 13; total 260 ants). In order to only measure internal infections, ant pools were washed to remove any conidiospores still loosely attached to the outer surface of their cuticles. This was done by vortexing them in 500 µl 0.05% Triton-X solution for 1 min and subsequently rinsing all ants individually with 100 µl 0.05% Triton-X solution.

### Quantification of super-infection load in micro-infected ants

The fungal load of the heterologous pathogen was determined using quantitative real-time PCR. Prior to DNA extraction, the samples were homogenized in a TissueLyser II (Qiagen) using a mixture of 2.8 mm ceramic (VWR), 1 mm zirconia (BioSpec Products) and 425-600 µm glass beads (Sigma-Aldrich). Homogenization was carried out in two steps (2× 2 min at 30Hz). DNA extractions were performed using Qiagen DNeasy96 Blood and Tissue Kit per the manufacturer’s instructions, with a final elution volume of 50 µl Buffer AE.

We then performed a real-time PCR assay to quantify the fungal ITS2 rRNA gene copies. Targeting this multi-copy gene ensures a high sensitivity and prevents any cross-amplification of the two fungi present in the ants. Quantification standards were obtained by extracting DNA of pure *Metarhizium* and *Beauveria* conidiospores. Site specific primers for *Beauveria bassiana* were taken from [69] (F: 5’- GCCGGCCCTGAAATG G; R: 5‘ - GATTCGAGGTCAACGTTCAGAA). Primers for the amplification of *Metarhizium robertsii* were designed based on GenBank sequence AY755505.1 (F: 5’- CCCTGTGGACTTGGTGTTG, R: 5’- GCTCCTGTTGCGAGTGTTTT), as in [70]. Both primer pairs were shown to not cross-amplify with the other fungal species.

Amplification was carried out in 20 µl reactions using 1x KAPA SYBR Fast qPCR master mix (KapaBiosystems), 3 pmol (*Metarhizium*) or 4 pmol (*Beauveria*) of each specific primer (Sigma-Aldrich) and 2 µl template on a CFX96 real-time PCR instrument (bio-rad). Cycling parameters were chosen according to manufacturer’s recommendations (annealing temperatures: 64 °C for *Metarhizium*, 60 °C for *Beauveria*). Quantification was based on a standard curve, with standards covering a range from 10^-2^ to 10^-5^ ng/µl fungal DNA for *Metarhizium* and 10^-2^ to 10^-4^ ng/µl fungal DNA for *Beauveria*. The respective lowest standard was determined to be the detection threshold. Each run included a negative control. Specificity was confirmed by performing a melting curve analysis after each run.

### Statistical analyses

All statistical analyses were carried out in the program R, version 3.3.2 [71], and all reported *P*-values are two-sided. To test for the overall effect of a homologous vs heterologous combination of micro-infection (with non-infected ants as a control) and secondary pathogen exposure, either via direct experimental exposure or social contact to a contaminated nestmate, we performed a global analysis with ‘treatment group’ (containing three levels, non-infected, homologous combinations and heterologous combinations) as a main effect into the respective models. To further test for the robustness of this overall pattern across the two pathogens and their exposure order, we performed an additional detailed analysis, in which we tested for the main effects of ‘micro-infection history’, which was split up into *Metarhizium* micro-infected, or *Beauveria* micro-infected, and non-infected controls, and ‘secondary pathogen exposure’ (direct exposure or contaminated nestmate) with either *Metarhizium* or *Beauveria*, as well as their interaction. As we reused data for the global and detailed statistical analysis, we corrected for multiple testing with the Benjamini-Hochberg correction and present adjusted p values.

To assess the significance of main effects of all models, we compared full models to null (intercept only) and reduced models (for those with multiple predictors), using likelihood ratio tests. We checked the appropriate diagnostics for all models, including over dispersion, Cook’s distance, dfbetas, dffits, leverage, variance inflation factors (package ‘car’, version 2.0-19 [72]), distribution of residuals, residuals plotted against fitted values and Levène’s test of equality of error variances – to test for obvious influential cases, outliers and deviations from the assumptions of normality and homogeneity of residuals. Where post hocs were necessary, we performed Tukey post hoc comparisons and adjusted the resulting *P*-values using the Benjamini-Hochberg procedure, using the ‘multcomp’ package (version 1.4-6 [73]).

To test for the effect of infection history on the survival of ants after homologous and heterologous pathogen challenge, we used Cox mixed effects models (package ‘coxme’, version 2.2-5 [74]), with ‘survival’ as the response variable. In the global analysis, treatment was the only main effect, but we included ‘replicate petri dish’ as a random intercept effect, since ants from the same petri dish are non-independent (Fig. 1). In the detailed analysis testing the two pathogens separately in their homologous/heterologous combinations (taking exposure order into account), we ran a Cox mixed effects models containing ‘infection history’, ‘nestmate contamination’ and their interaction as fixed effects. As the interaction was significant, we performed individual models comparing the survival of micro-infected ants to non-infected control ants following a pathogen challenge separately, by including ‘infection history’ (non-infected *versus* micro-infected) as a fixed factor (Fig. S1). Again, ‘replicate petri dish’ was included as a random effect. We controlled for multiple testing by adjusting *P* values using the Benjamini-Hochberg procedure to protect against a false discovery rate of 0.05.

When testing for the effect of pathogen co-infection after direct co-exposure (Fig. S2), survival of naive ants after exposure with either *Metarhizium* or *Beauveria*, or a simultaneous co-exposure with both fungal pathogens, we used a Cox proportional hazards regression (package ‘survival’, version 2.38 [75]). As the overall regression model was significant we performed post hoc comparisons to test for differences between the respective treatments.

General(ised) linear models (GLM) were used to analyse whether the behaviour of ants – aggression, poison spraying and allogrooming – depended on their micro-infection history and contamination treatment of nestmates, both performing the global (Fig. 2) and detailed analysis (Fig. S3). For the grooming data, we used GLMs with Gaussian errors. Since the aggression and poison spraying behavioural data followed a negative binomial distribution, we analysed this data using generalised linear models with negative binomial errors (package ‘MASS’, version 7.3-47 [76]). For the detailed analyses, we also ran separate models for each nestmate ‘treatment’ (contamination with *Metarhizium-*, or *Beauveria*) with ‘infection history’ as a sole predictor, on which we performed post hocs to obtain comparisons between the different types of ‘infection history’ (non-infected, *Metarhizium*, *Beauveria*). For the detailed analysis of aggression, the interaction between micro-infection history and nestmate contamination was non-significant (GLM, LR χ^2^ = 3.9, df = 2, *P* = 0.142), so it was removed and the model re-run without it, to obtain better estimates of the remaining predictors. In addition, as there was no poison spraying by non-infected ants towards *Beauveria*-contagious nestmates, we artificially added a single spraying event to this group to gain finite estimates and avoid complete separation of the data. Our model thus overestimates spraying behaviour in non-infected ants, indicating that the already significant differences between the different types of ‘infection history’ in reality are even greater.

The transfer of infectious conidiospores from contaminated individuals to micro- infected nestmates was analysed using GLMs with Gaussian errors after data were ln(x+1) transformed to fulfil the assumption of normality. Again, we performed both a global analysis (Fig. 3A) and detailed analysis for the two pathogens (Fig. S4). The super-infection load of micro-infected ants was analysed using linear mixed effects regressions (LMER), with presence/absence of behavioural change (risk-adjusted, control) included as predictor for the global analysis (Fig. 3B). For the detailed analysis, we also included ‘micro infection history’ as a second predictor, and performed separate LMERs to determine how super-infection loads changes within both pathogen species (Fig. S5). We square root transformed infection load to achieve normality. Since the DNA extraction was carried out over two separate runs, ‘run’ was included as a random intercept effect into all models. Three different experimenters performed the pathogen exposure, so a random intercept was also included for ‘person’. From our 52 original samples, we excluded one outlier that showed exceptionally high fungal multiplication, reaching a 250-fold higher value than the maximum value of all other samples.

## Acknowledgements

We thank B.M. Steinwender and J. Eilenberg for the fungal strains and the whole *Social Immunity* Team at IST Austria for support and discussion throughout the project, particularly B. Milutinović, M.A. Fürst, B. Casillas-Pérez and M. Iglesias. We also thank P. Schmid-Hempel, J. Kurtz, S.A. Frank, C.D. Nunn, M. Schaller, C. Russell and F. Jephcott for discussion, and R.M. Bush, L.V. Ugelvig and M. Sixt, for comments on the manuscript. Funding was obtained by the European Research Council (Starting Grant 240371 to S.C.).

